# Assessing whitefly diversity to infer about begomovirus dynamics in cassava in Brazil

**DOI:** 10.1101/2020.09.22.309187

**Authors:** César A.D. Xavier, Angélica M. Nogueira, Vinícius H. Bello, Luís F. M. Watanabe, Miguel Alves-Júnior, Leonardo F. Barbosa, José E.A. Beserra-Junior, Alessandra J. Boari, Renata F. Calegario, Eduardo S. Gorayeb, Jaime Honorato-Júnior, Gabriel Koch, Gaus S.A. Lima, Cristian A. Lopes, Raquel N. Mello, Késsia F. C. Pantoja, Fabio N. Silva, Roberto Ramos-Sobrinho, Enilton N. Santana, José W.P. Silva, Renate Krause-Sakate, F.M. Zerbini

## Abstract

Plant virus ecology is strongly dependent on that of its vector. The necessity of a competent vector for transmission is a primary ecological factor driving the host range expansion of plant arthropod-borne viruses, with vectors playing an essential role in promoting disease emergence. Cassava begomoviruses severely constrain cassava production in Africa. Curiously, begomoviruses have never been reported in cassava in South America, the center of origin for this crop. It has been hypothesized that the absence of a competent begomoviruses vector that efficiently colonizes cassava is the reason why begomoviruses have not emerged in South America. To test this hypothesis, we performed a country-wide whitefly diversity study in cassava in Brazil. Adults and/or nymphs of whiteflies were collected from sixty-six cassava fields across twelve states representing the main agroecological zones of the country. A total of 1,385 individuals were genotyped based on partial mitochondrial cytochrome oxidase I (mtCOI) sequences. A high species richness was observed, with five previously described species and two putative new ones. The most prevalent species were *Tetraleurodes acaciae* and *Bemisia tuberculata*, representing over 75% of the analyzed individuals. Although we detected, for the first time, the presence of *Bemisia tabaci* Middle East-Asia Minor 1 (*Bt*MEAM1) colonizing cassava in Brazil, it was not prevalent. The species composition varied across regions, with fields in the Northeast region showing a higher diversity. These results expand our knowledge of whitefly diversity in cassava and support the hypothesis that begomovirus epidemics have not occurred in cassava in Brazil due to the absence of competent vector populations. However, they indicate an ongoing adaptation process of *Bt*MEAM1 to cassava, increasing the likelihood of begomovirus emergence in this crop in the near future.

## Introduction

Cassava (*Manihot esculenta* Crantz) is a perennial shrub of the Euphorbiaceae family with great economic and social importance, especially in Africa, Asia, and Latin America. Currently, cassava is the third most important source of calories after rice and corn and is a staple food for more than one billion of people living mainly in developing countries (1). Although the botanical and geographical origin of *M. esculenta* is still debated, studies based on genetic markers and archaeological evidence suggest that domesticated cassava originated from the wild relative progenitor *M. esculenta* ssp. *flabellifolia* in the Amazon basin, with the domestication center located at the southern border of the Amazon in Brazil (2-5). After its introduction in west Africa by Portuguese traders during the 16^th^ century, cassava quickly disseminated throughout tropical Africa and Asia (6). Currently, the African continent is the world’s biggest cassava producer, followed by Asia and South America (FAO, 2017). Due to its high resilience to adverse environmental conditions, especially drought, high yield per unit of land and low level of management and inputs required during its life cycle, cassava is a suitable crop for poor and small farmers, partially ensuring food security in many African countries (7-9).

Nevertheless, cassava may be affected by several pathogens and pests. Whiteflies (Hemiptera: Aleyrodidae) are one of the major constraints to its production in developing countries (10). Whiteflies comprise a diverse group of phloem-feeding insects, with more than 1,500 species assigned to 126 genera of which over 20 species have been reported to colonize cassava worldwide (11). In addition to the direct damage due feeding in the plant phloem, whiteflies cause indirect damage by deposition of honeydew, favoring the growth of sooty mold fungi on the leaf surface, and mainly by transmission of a broad range of viruses (12). Currently, species included in the genera *Aleurodicus, Aleurothrixus, Bemisia* and *Trialeurodes* have been shown to constitute effective vectors of plant viruses classified in five families (12-15). *Aleurodicus dispersus* and *Aleurothrixus trachoides* each transmit only one virus from the genera *Ipomovirus* and *Begomovirus*, respectively, while *Trialeurodes vaporariorum* and *T. abutilonea* transmit a few viruses included in the genera *Crinivirus* and *Torradovirus*. On the other hand, the *Bemisia tabaci* complex comprises one of the most important group of plant virus vectors, transmitting over 430 viruses, the majority included in the genus *Begomovirus* (12, 16) but including also viruses classified in the genera *Carlavirus, Crinivirus, Ipomovirus, Polerovrius* and *Torradovirus* (12, 17-20).

Over the last decade, advances in the use of molecular markers has led to a deep reappraisal of the taxonomic status of *B. tabaci* (21, 22). Based on molecular phylogeny of the mitochondrial cytochrome oxidase I (mtCOI) gene, it has been proposed that *B. tabaci* consists of a complex of more than 40 cryptic (morphologically indistinguishable) species (21-25). Partial or complete reproductive isolation and biological and ecological differences among distinct species within the complex support the proposed classification (18, 26-28). The global dissemination of polyphagous and invasive species, such as *B. tabaci* Middle East-Asia Minor 1 (*Bt*MEAM1) and *B. tabaci* Mediterranean (*Bt*MED), have caused major changes in the epidemiology of crop-infecting begomoviruses such as *Tomato yellow leaf curl virus* (TYLCV), currently present in all the main tomato producing areas of the world (29-31). In addition, the dissemination of polyphagous whiteflies has favored the transfer of indigenous begomoviruses from wild reservoir hosts to cultivated plants, as occurred in tomato crops in Brazil after the introduction of *Bt*MEAM1 in the mid-1990’s (32, 33).

Specific associations between endemic populations of *B. tabaci* and indigenous begomoviruses have also led to the emergence of severe epidemic in crops (34, 35). Cassava mosaic disease (CMD) is considered the most significant constraint to cassava production in Africa (36, 37). CMD is caused by viruses of the genus *Begomovirus* (family *Geminiviridae*), which are transmitted in a circulative manner by whiteflies of the *Bemisia tabaci* cryptic species complex (16). To date, nine cassava mosaic begomoviruses (CMBs) have been reported in association with CMD, seven of them in Africa and two in the Indian subcontinent (37-39). The emergence of CMD seems to have been the result of the transfer of indigenous begomoviruses from wild reservoir hosts to cassava, probably mediated by endemic populations of *B. tabaci* that have adapted to feed in cassava since its introduction from South America (34, 40). Although wild reservoir hosts and a possible ancestral progenitor of current begomoviruses causing CMD have not been found, the absence of cassava-infecting begomoviruses in the Americas supports an African origin for those viruses, and the presence of cassava-adapted *B. tabaci* species being restricted to Africa reinforces that hypothesis. The high species diversity and high level of molecular variation observed in viral populations causing CMD strongly suggests Africa as a diversification center for CMBs, with distinct CMBs recurrently emerging and evolving for a long time (41-44).

Even if CMD in Africa is caused by indigenous viruses, the fact that cassava in South America has not been affected by begomoviruses is puzzling (38, 45, 46). Carabali, Bellotti (47) suggested that the absence of cassava-infecting begomoviruses in the Americas would be due to lack of competent *B. tabaci* species that efficiently colonize cassava (34, 40). In Colombia, the inability of *B. tabaci* MEAM1 to colonize *M. esculenta* efficiently has been demonstrated under experimental conditions, reinforcing the above hypothesis (46). In addition, a recent study also from Colombia failed to detect any whitefly species of the *B. tabaci* complex in cassava (48).

Although whitefly diversity in Brazil has been surveyed extensively in recent years (49-54), no study has been carried out specifically to explore the composition of whitefly communities colonizing cassava. Those studies carried out in other crops demonstrated that *B. tabaci* MEAM1 is the predominant species across Brazil in crops such as common bean, cotton, pepper, tomato and soybean. Furthermore, *B. tabaci* MED, which was recently introduced in Brazil, has quickly spread and currently is present in five states from the South and Southeast regions (52-54). A small number of whitefly samples from cassava were analyzed in those studies, with *B. tuberculata* and *Tetraleurodes acaciae* prevalent and detected exclusively in cassava (51, 54). A large survey addressing whitefly diversity in cassava in its domestication center could provide clues to understand the absence of a CMD-like disease in the Americas. Moreover, this knowledge would be useful to anticipate the potential of emergence of begomoviruses in the crop and to help anticipate a management strategy.

Given this context, the objective of this work was to evaluate whitefly diversity in cassava across Brazil to infer about the absence of begomovirus epidemic in cassava. Our results demonstrated that the most prevalent species in cassava were *T. acaciae* and *B. tuberculata*. In addition, we detected for the first time the presence of *Bt*MEAM1 colonizing cassava in Brazil. The possible implications of these findings are discussed considering the absence of CMD and the potential for its emergence in cassava fields in Brazil.

## Methods

### Whitefly and cassava samples

Whiteflies were collected exclusively from cassava (*M. esculenta*) plants across 12 Brazilian states representative of the five macroregions (North, Northeast, Midwest, Southeast and South; Figure 1) between March 2016 and February 2019 (Table 1). To gather evidence of whether a given species was colonizing cassava, adults and nymphs from the same field were collected whenever possible (Table 1). Samples were obtained from commercial and non-commercial (subsistence) crops. Whitefly adults were sampled using a hand-held aspirator and nymphs were collected with the aid of a needle. Insects were preserved in 95% ethanol and stored at −20°C until being used for molecular identification of the species.

**Table 1.**
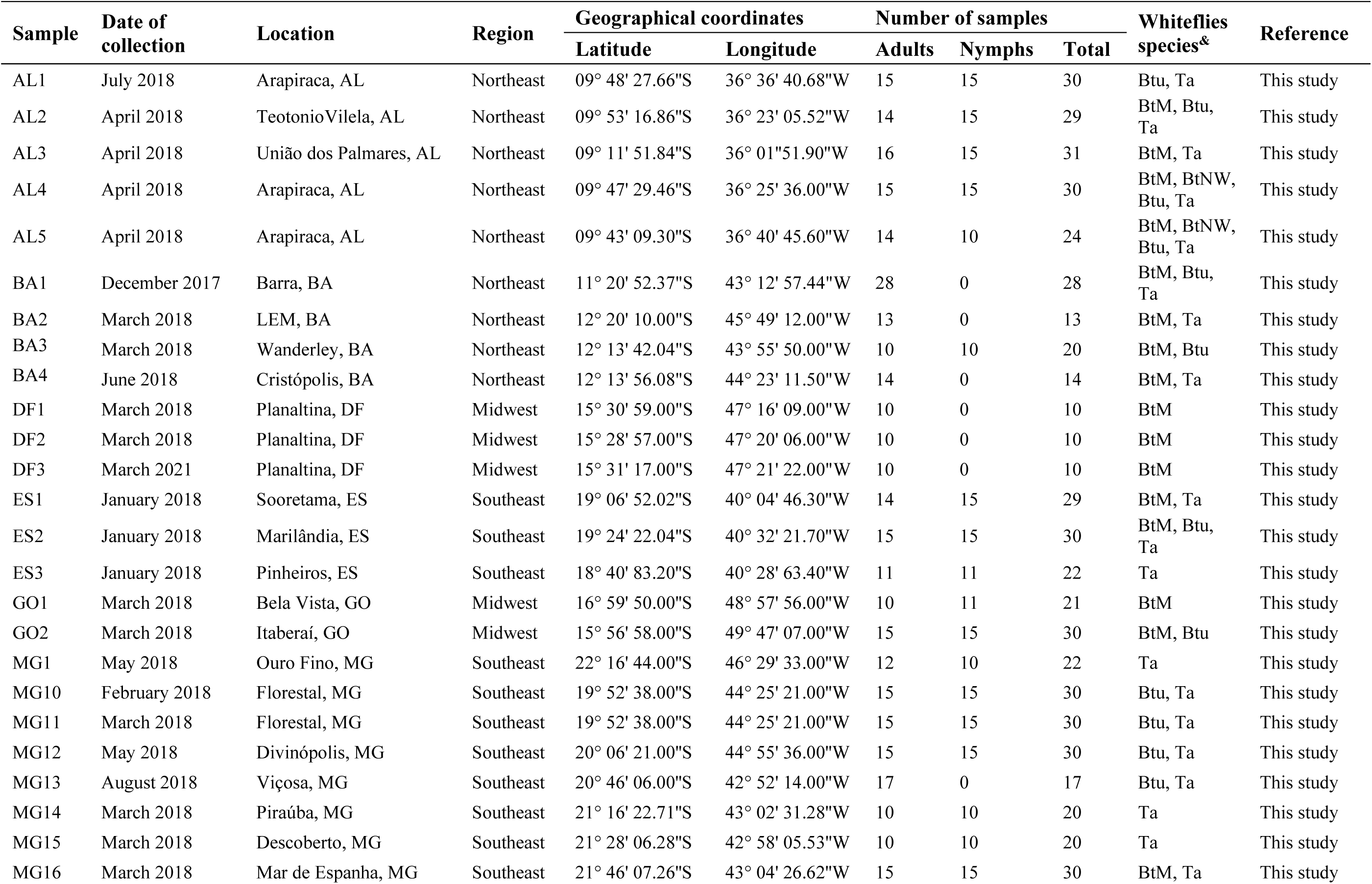
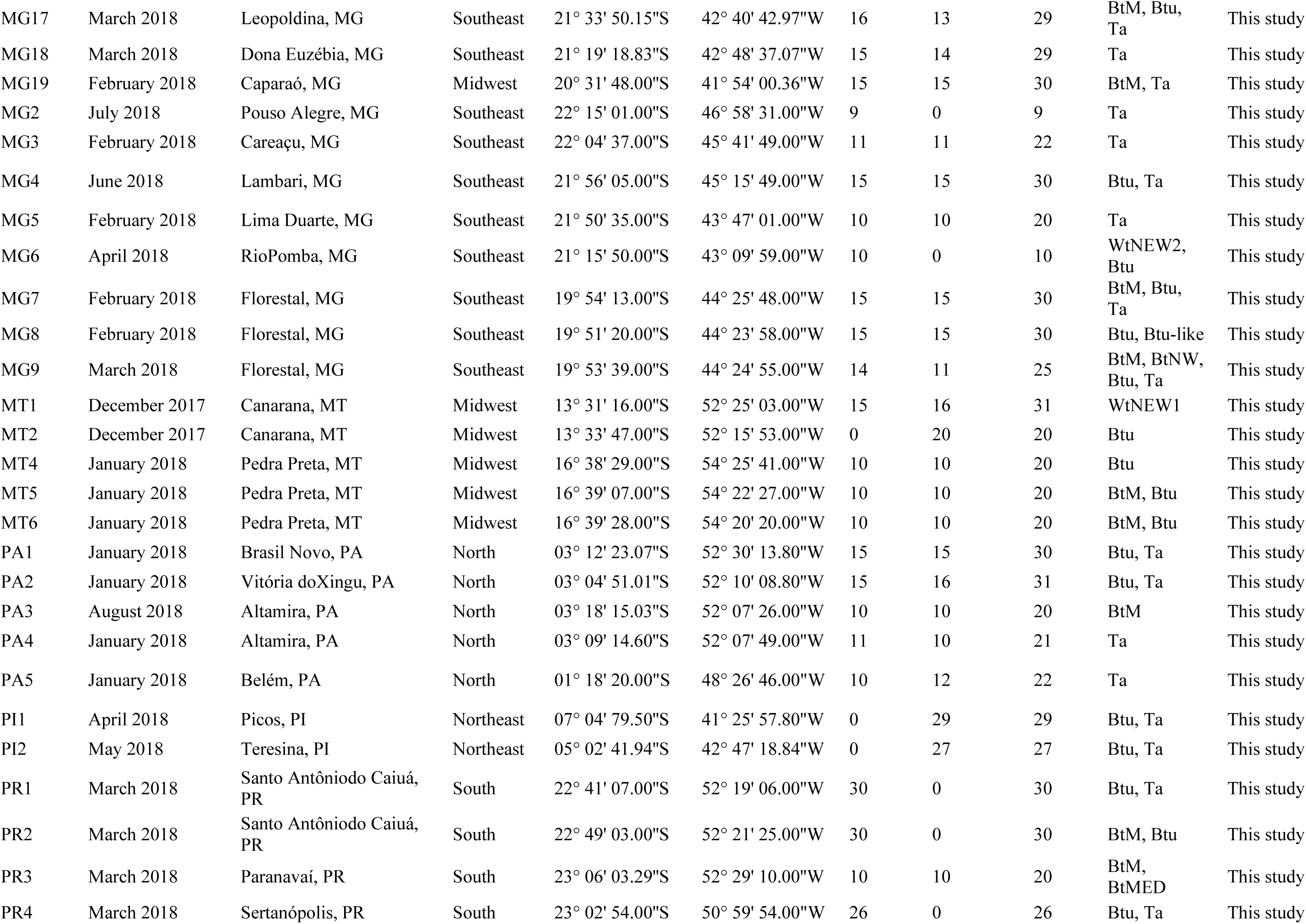
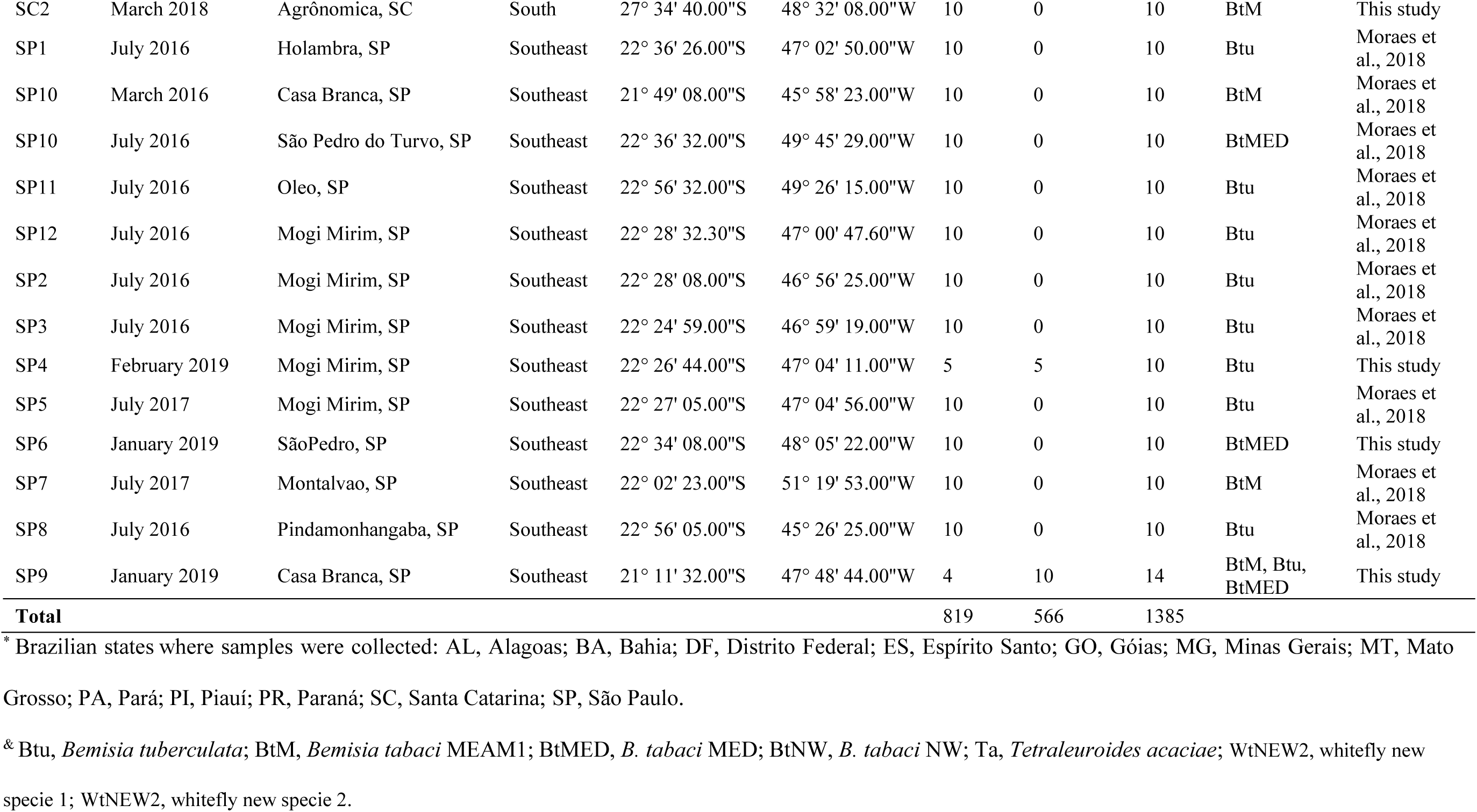
Sampled sites and whiteflies species detected in cassava in Brazil.

**Figure 1.**
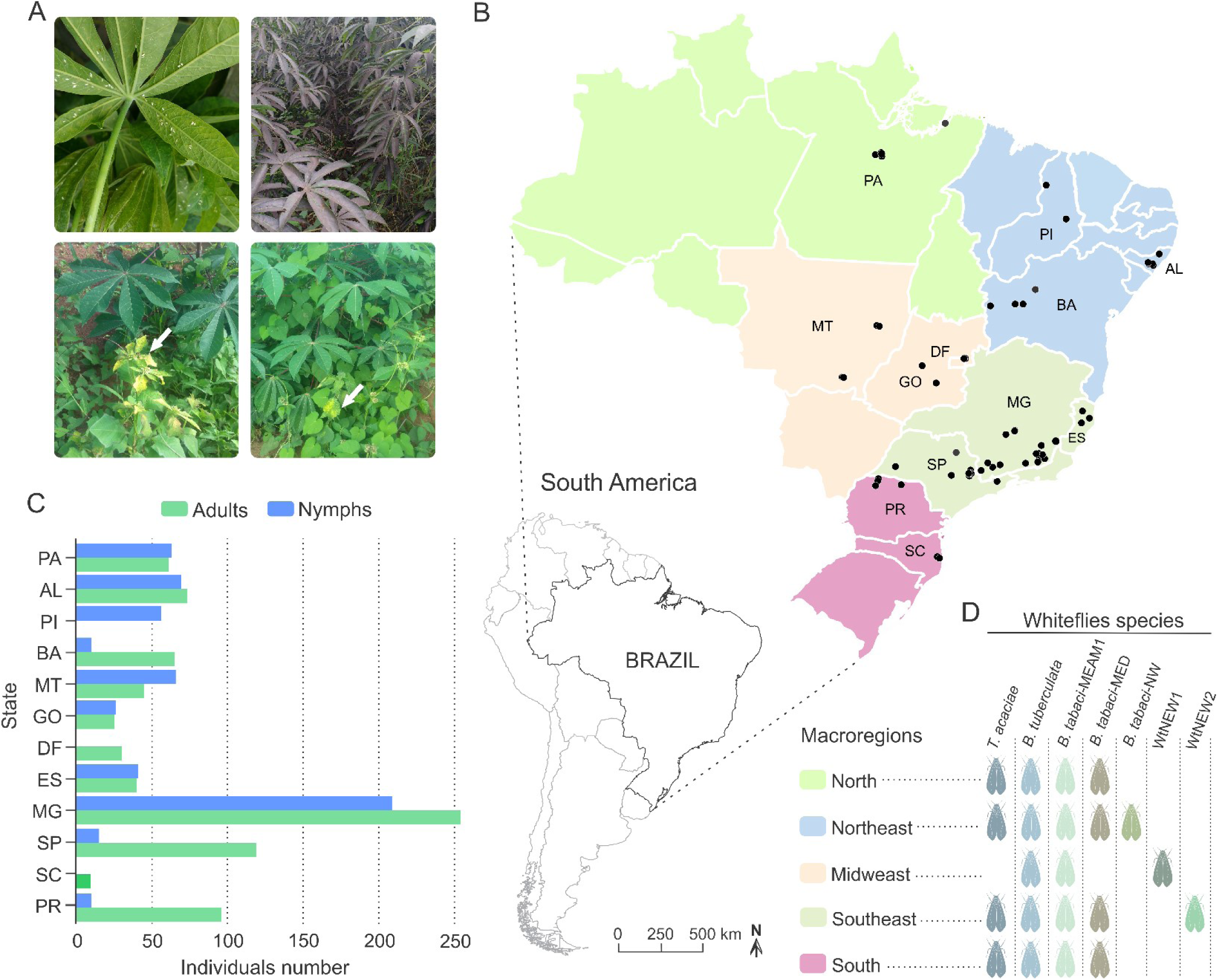
**A**. Clockwise from top-left: Adults and nymphs of *Bemisia tuberculata* colonizing cassava in Mogi Mirim, São Paulo state. Growth of sooty mould fungus on the leaf surface due to the deposition of honeydew by whiteflies. Presence of begomovirus-infected *Blainvillea rhomboidea* (family Asteraceae) in a cassava field in Minas Gerais state. Presence of begomovirus-infected *Euphorbia heterophylla* (family Euphorbiaceae) in a cassava field in Minas Gerais state. **B**. Map of Brazil showing the locations where whiteflies samples were collected. The map is colored according to the regions as indicated in the legend. Blacks dots correspond to the sampled sites. Scale bar is only for Brazil map. **C**. Number of adults and nymphs analyzed from each sampled site according to state. **D**. Specie distribution according to region. AL, Alagoas; BA, Bahia; DF, Distrito Federal; ES, Espírito Santo; GO, Goiás; MG, Minas Gerais; MT, Mato Grosso; PA, Pará; Piauí; PR, Paraná; SC, Santa Catarina; SP, São Paulo.

To verify the presence of begomoviruses infecting cassava, foliar samples were also collected at some sampled sites (Suppl. Table S1). The samples were collected randomly regardless of the presence of virus-like symptoms. The leaves were press-dried and stored at room temperature as herbarium-like samples until being used for DNA extraction.

### Whitefly species identification

Whitefly species were identified using PCR-RFLP of the partial mtCOI fragment followed by sequencing, as previously described (54). When enough adults and nymphs were collected at a given sampled site, ten individuals from each stage were analyzed, and when only one stage was obtained, 20 individuals were tested (Table 1). When variation in the RFLP pattern was observed in the first screening, suggesting that more than one species could be present in that site, approximately five additional individuals for each stage were analyzed according to sample availability.

Total DNA was extracted from single individual whiteflies following a Chelex protocol (55). Briefly, adults or nymphs were ground in 30 µl of Chelex buffer (5% Chelex in 1x Tris-EDTA) using a toothpick in a 600 µl tube. Samples were vortexed for 30 seconds and incubated at 99°C for 8 min in a PTC-100 thermocycler (MJ Research). Next, the tubes were centrifuged at 14,000 g for 5 min and 20 µl of the supernatant was collected and transferred to a new tube. One microliter of the supernatant was used as a template for PCR amplification of a 800 bp fragment of the mtCOI gene using primers C1-J-2195 and L2-N-3014 (56, 57). PCR was performed using 0.2 µM of forward and reverse primers in a final volume of 25 µl using GoTaq Colorless Master Mix (Promega), following the manufacturer’s instructions. The PCR cycles consisted of an initial denaturing step at 95°C for 5 min, followed by 35 cycles at 95°C for 30 sec, 42°C for 45 sec and 72°C for 1 min, with a final extension at 72°C for 10 min. Amplified products were visualized in 0.8% agarose gels stained with ethidium bromide and directly used for RFLP analysis (58).

RFLP analysis of the amplicons consisted of 5 µl of each PCR product digested with 0.1 unit of *Taq*I (Promega) in a final volume of 20 µl. Reactions were performed at 65°C for 2 hours and visualized in 1.2% agarose gels stained with ethidium bromide. To verify whether the predicted mtCOI restriction pattern corresponded to a given species according to *in silico* prediction, a subset of PCR products from adults and nymphs representative of distinct patterns from different sampled sites were selected and sequenced. PCR products were precipitated with 100% ethanol and 3 M sodium acetate pH 5.2 (59) and sequenced commercially (Macrogen Inc.) in both directions using primers C1-J-2195/L2-N-3014.

For a small subset of samples that failed to yield a PCR product using primers C1-J-2195 and L2-N-3014, a second screening, using a recently described primer set with improved specificity for species of the *B. tabaci* complex and *B. afer* (2195Bt and C012/Bt-sh2), was performed (24). Samples that still failed to amplify or had unexpected RFLP pattern were analyzed with specific primers for *T. vaporariorum* (TvapF and Wfrev) (60).

### Sequence comparisons and phylogenetic analysis

Nucleotide sequences were first checked for quality and assembled using Geneious v. 8.1 (61). mtCOI sequences were initially analyzed with the BLAST*n* algorithm (62) to determine the whitefly species with which they shared greatest similarity. Pairwise comparisons between all mtCOI sequences obtained here and those with higher similarities (as determined by the BLAST*n* search) were performed with the program SDT v. 1.2 (63) using the MUSCLE alignment option (64).

For phylogenetic analyses, the final dataset was composed of 142 sequences: 95 obtained in this work and 47 sequences representative of species in the family Aleyrodidae. Sequences were retrieved from GenBank and from the updated mtCOI reference dataset for species of the *Bemisia tabaci* complex (65). Multiple sequence alignments were prepared using the MUSCLE option in MEGA7 (66). Alignments were checked and manually adjusted when necessary. Phylogenetic trees were constructed using Bayesian inference performed with MrBayes v. 3.0b4 (67). The program MrModeltest v. 2.2 (68) was used to select the nucleotide substitution model with the best fit in the Akaike Information Criterion (AIC). The analyses were carried out running 50,000,000 generations with sampling at every 1,000 generations and a burn-in of 25%. The convergence was assumed when average standard deviation of split frequencies was lower than 0.001. Trees were visualized and edited using FigTree (tree.bio.ed.ac.uk/software/figtree) and CorelDRAW X5, respectively.

### Virus detection in foliar samples

Total DNA was extracted as described (69) and used as a template for PCR using the DNA-A universal primer pair PAL1v1978 and PAR1c496 (70). PCR was performed in a final volume of 25 µl using *Taq* DNA Polymerase (Invitrogen) following the manufacturer’s instructions. The PCR cycles consisted of an initial denaturing step at 95°C for 5 min, followed by 35 cycles at 95°C for 1 min, 52°C for 1 min and 72°C for 1 min, with a final extension at 72°C for 10 min. PCR products were visualized in 0.8% agarose gels stained with ethidium bromide

### Diversity index and statical analysis

Simpson’s index of diversity (1-D) was calculated to verify if there was any difference in whitefly diversity across macroregions. This index represents the probability that two randomly chosen individuals in a given sampled site will belong to distinct species (71). Simpson’s index was chosen as its value increases with increasing diversity and assigns more weight to more abundant species in a sample. We assume that species colonizing cassava will be in abundance, whereas rare species that briefly visit the plant without colonizing it will be underrepresented. Simpson’s index was calculated for each sampled site separately and then pooled according to macroregions. To assess the statistical significance of the differences in diversity among regions, the non-parametric Kruskal-Wallis test followed by *post hoc* multiple comparison test using Fisher’s least significant difference was calculated, using the function kruskal implemented in the Agricolae package in R software (72). Non-parametric Spearman’s rank correlation coefficient analysis was performed using the ggpubr package in R software (72).

## Results

### High whitefly species richness in cassava in Brazil

To verify the composition of whitefly communities colonizing cassava in Brazil, sampling was performed across the country, including the main agroecological zones. A total of 66 sites from 12 states were sampled (Figure 1; Table 1). Out of 1,385 individuals submitted to PCR-RFLP analysis, 58 adults and 37 nymphs from different locations and representing distinct restriction patterns were sequenced. The combination of PCR-RFLP followed by sequencing showed reliability and consistence for species identification without misidentification due to incongruence between the two methods.

Based on pairwise comparisons and molecular phylogeny of the partial mtCOI gene, we identified the presence of at least seven species comprising the whitefly community in cassava (Figure 2; Table 1). Among them, *T. acaciae* and *B. tuberculata*, both previously reported in this crop, were the most prevalent, representing over 75% of the analyzed individuals. In addition, based on the criterion of 3.5% divergence to differentiate species within the *B. tabaci* complex, three *B. tabaci* species were identified, with *Bt*MED previously reported, and *Bt*MEAM1 and *Bt*NW identified for the first time in cassava fields in Brazil (Figure 2; Table 1). The species *Bt*MEAM1 represented 18% of the total individuals analyzed, followed by *Bt*MED (1.6%) and *Bt*NW (0.21%).

**Figure 2.**
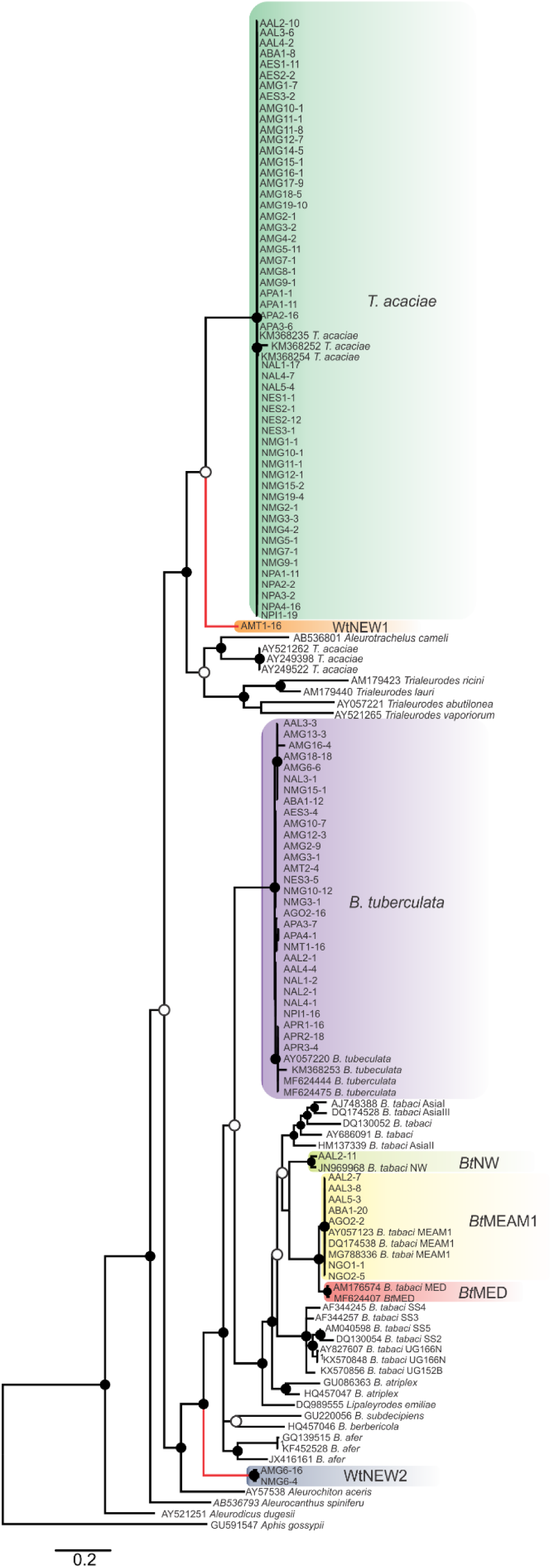
Bayesian phylogenetic tree based on partial nucleotide sequences of the mitochondrial cytochrome oxidase (mtCOI) gene of representative individuals of each whitefly species detected in this study and reference sequences retrieved from GenBank. The tree was rooted with the aphid *Aphis gossypii*. Bayesian posterior probabilities are shown at the nodes. The scale bar represents the number of nucleotide substitutions per site. Nodes with posterior probability values between 0.60 and 0.80 are indicated by empty circles and nodes with values equal to or greater than 0.81 are indicated by filled circles. Clades highlighted with different colors indicate the species detected in this study. Branches highlighted in red indicate the putative new species detected here.

Furthermore, two putative new species were identified (Figure 2), provisionally named whitefly new species 1 and 2 (WtNEW1 and WtNEW2). The WtNEW1 mtCOI sequence (KY249522) showed highest identity (80.65%) and clustered close to the *T. acaciae* clade, comprised of individuals reported here and three other previously reported sequences from cassava in Brazil (Figure 2). For WtNEW2, two mtCOI sequences obtained from an adult (JX678666) and a nymph (DQ989531) shared 97.81% among them and showed highest identity with *B. tabaci* (adult: 82.11%; nymph: 81.68%) and clustered as a basal sister clade to the genus *Bemisia* (Figure 2). Although whitefly taxonomy is predominantly based on puparial characters (73) and there is no taxonomic criterion established based in mtCOI sequences for most of the groups, as has been proposed for the *B. tabaci* complex, the level of divergence between the two proposed new species with the closest species is similar to the level of divergence observed between species already described within the Aleyrodidae, as demonstrated in pairwise comparisons (data not shown) and phylogenetic analysis (Figure 2). Nevertheless, further molecular and morphological characterization should be performed. Together, these results indicate the existence of a high whitefly species richness in cassava in Brazil.

### Both the prevalence and the capacity to colonize cassava differ among species

Nymphs were collected for samples identified as *T. acaciae, B. tuberculata, Bt*MEAM1 and the two new putative species (Figure 3A), suggesting that these species may colonize cassava. Nymphs were not obtained at two sites where *Bt*MED was prevalent (SP1 and SP12). Although it could be suggested that this species has the potential to colonize cassava due to the high prevalence of adults at these two sites, the lack of nymphs suggests otherwise. Moreover, at the sites PR4 and MT6, *Bt*MEAM1 predominated among adults but 100% of the nymphs were *B. tuberculata*, suggesting that the predominance at one stage does not necessarily mean predominance in another stage. Indeed, correlation analysis between the number of adults and nymphs, performed for all sites where both stages were sampled, showed no significant correlation between them (Supp. Figure S1). Further sampling in those sites or free-choice experiments are necessary to confirm the potential of *Bt*MED to colonize cassava. Considering the whole sampling, we detected only three adults of *Bt*NW, suggesting an inability of this specie to colonize cassava.

**Figure 3.**
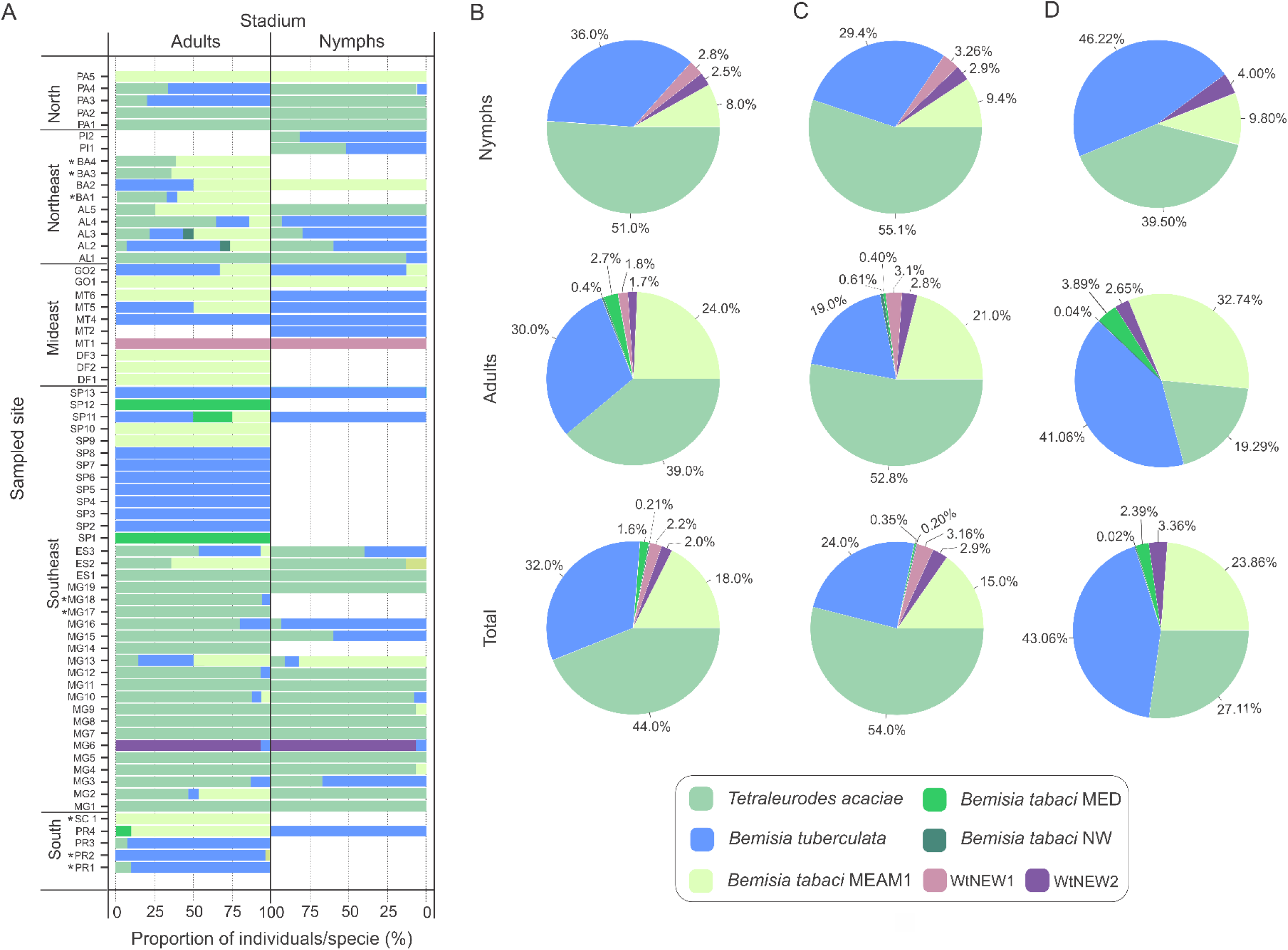
Composition of whitefly populations colonizing cassava in Brazil. **A**. Species composition at each sampled site according to stage of development (adult and nymphs). Asterisks indicate that nymphs were not detected. **B, C, D**. Species distribution of the 1,385 individuals genotyped in this study considering the samples from all sites (**B**) or only samples from sites where both adults and nymphs were sampled (**C**) or without samples from Minas Gerais state (**D**).

To verify if prevalence differs among species across distinct developmental stages, the data were separated according to stage and the proportions of individuals were compared for the three most abundant species (Figure 3B, C). Considering the entire data set, *T. acaciae* was the prevalent species, followed by *B. tuberculata* and *Bt*MEAM1 (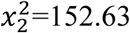, *P*<2.2×10^−16^). The same was true according to stage, either adults (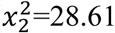, *P*<6.1×10^−07^) or nymphs (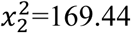, *P*<2.2×10^−16^; Figure 3B). However, caution is needed to interpret these results as only adults were sampled at some sites where *Bt*MEAM1 and *B. tuberculata* were prevalent (Figure 3A), which could bias the analysis, causing an underestimation of the number of nymphs for those species. Therefore, we also analyzed the data considering only those sites where both adults and nymphs were obtained. Again, *T. acaciae* was the predominant species followed by *B. tuberculata* and *Bt*MEAM1 considering either the entire data set (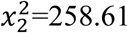, *P*<2.2×10^−16^) or only nymphs (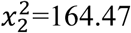, *P*<2.2×10^−16^). When only adults were considered, *T. acaciae* was still predominant (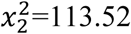, *P*<2.2×10^−16^) but no difference between *B. tuberculata* and *Bt*MEAM1 was observed (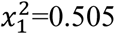,*P*=0.477; Figure 3C). Moreover, it could be argued that samples from Minas Gerais (MG) were overrepresented in our sampling (Figure 1C), which could also bias the results presented above due to the predominance of *T. acaciae* in this state (Figure 3A). To test this possibility, we analyzed the data excluding the samples from MG. In this case, when both stages were considered, *B. tuberculata* was predominant (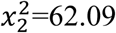, *P*=3.3×10^−14^) but no difference between *T. acaciae* and *Bt*MEAM1 was observed (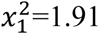, *P*=0.166). When each stage was considered separately, *B. tuberculata* was predominant followed by *Bt*MEAM1 and *T. acaciae* (adults: 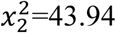, *P*=2.9×10^−10^; nymphs: 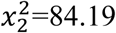, *P*<2.2×10^−16^). Together, these results indicate that the potential to colonize cassava differs among species, which could be due either to lower preference for the plant or to differences in the competitive ability among species during cassava colonization. In addition, they reinforce the low efficiency of *Bt*MEAM1 to colonize cassava.

### Competitive interference does not explain the differences in prevalence

Interestingly, at least two species were detected co-occurring at 51% of the sampled sites (Figure 3A). To verify the possibility of competition among *T. acaciae, B. tuberculata* and *Bt*MEAM1 to explain the observed differences in prevalence (instead of differences in host preference), the competitive capacity of these three species was inferred based on the analysis of predominance at the sites where they occurred together. Initially, we verified if there were any differences in incidence, defined here as the number of sampled sites where at least one individual belonging to one of the three species was detected (Figure 4A). The results demonstrate that there were no differences in incidence among them (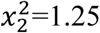, *P*=0.537; Figure 4A). In addition, no differences were observed when the proportion of sites where whitefly species occurred alone or in different combinations was compared (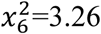, *P*=0.776; Figure 4B). However, when we compared the occurrence between *Bt*MEAM1 and non-*B. tabaci* species at the sites where they occur alone, the number of sites with non-*B. tabaci* species was higher (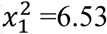, *P*=0.011; Figure 4B). Thus, the competitive capacity was inferred based on the proportion of individuals from each species at the fields where these species were detected co-occurring in different combinations (Figure 4C). Interestingly, at the sites where *Bt*MEAM1 and *B. tuberculata* were sampled together, *B. tuberculata* predominated over *Bt*MEAM1, suggesting higher competitive potential (Figure 4C). For all other species combinations, no evidence of differences in competitive capacity were observed (Figure 4C). Together, these results suggest that, rather than competition, lower host preference by *Bt*MEAM1 explains its non-prevalence compared to *T. acaciae* and *B. tuberculata*, resulting in low colonization rate as indicated by the low number of *Bt*MEAM1 nymphs detected in cassava (Figure 3).

**Figure 4.**
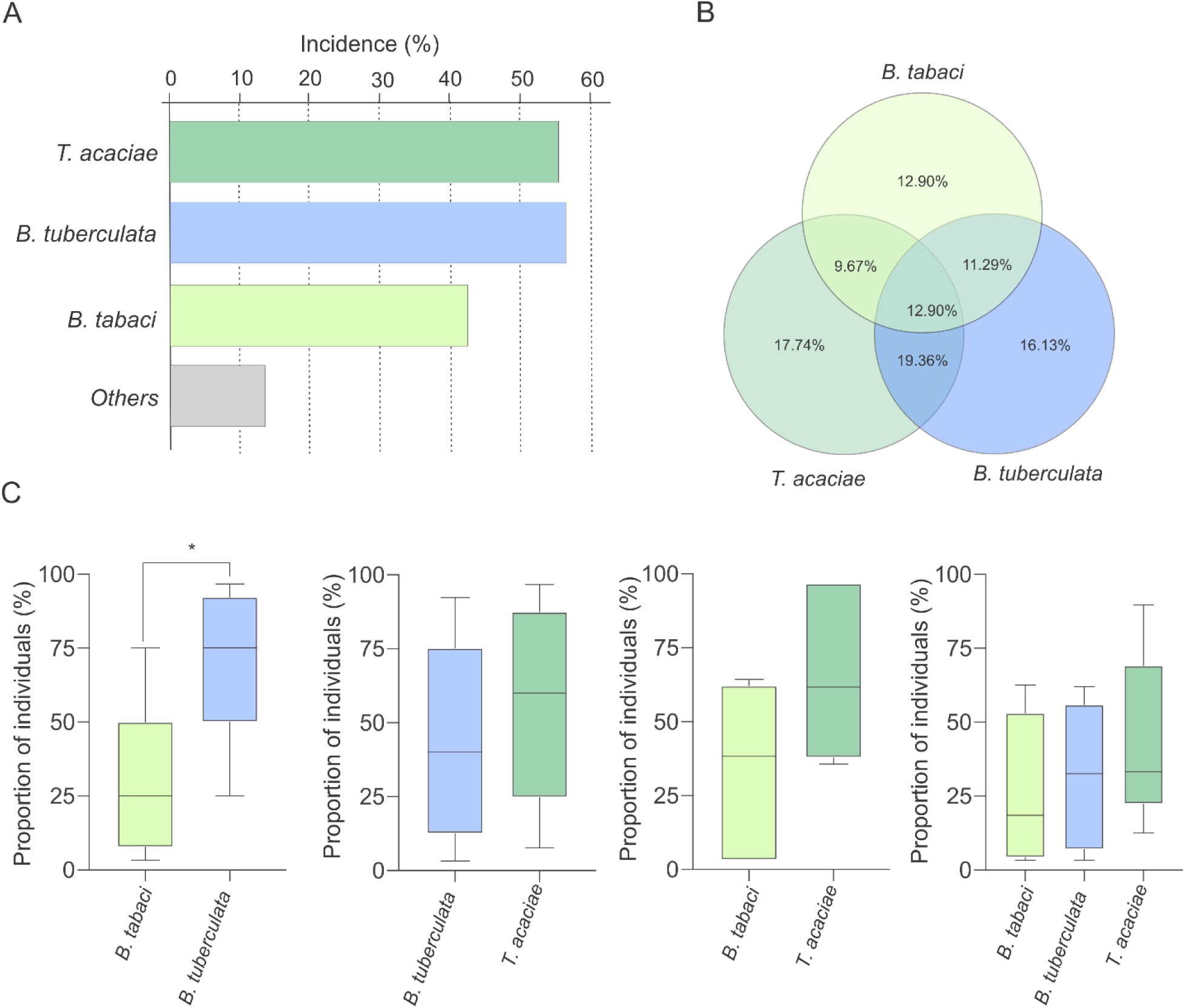
**A**. Incidence of *Trialeurodes acaciae, Bemisia tuberculata* and *B. tabaci* MEAM1 in cassava fields in Brazil, measured as the percentage of sampled sites where at least one individual belonging to each one of the three species was detected. Other species detected at low incidence are shown together as “others”. **B**. Venn diagram showing the proportion of sites where each one of the three whitefly species occur alone or in different combinations. **C**. Competitive capacity inferred based on the prevalence of individuals from each of two species in fields where those two species were detected co-occurring. The horizontal line inside the box corresponds to the median. The asterisk indicates a significant difference according to the non-parametric Kruskal-Wallis test (*p*<0.05).

### Composition and species diversity of whiteflies differ among Brazilian regions

The predominance of species composing the whitefly community across macroregions varied considerably. While *T. acaciae* predominated in the North, Southeast and Northeast, it was not detected in the Midwest (Figure 5A). In addition, *B. tuberculata* was detected in all regions, and was prevalent in the South and Midwest. *Bt*MEAM1, although not prevalent in any of the regions, was also detected in all regions. Although the number of species detected was higher in the Southeast, where six species out seven were detected, whitefly diversity was significantly higher in fields in the Northeast according to Simpson’s index of diversity (Figure 5B), with no differences among the other four regions.

**Figure 5.**
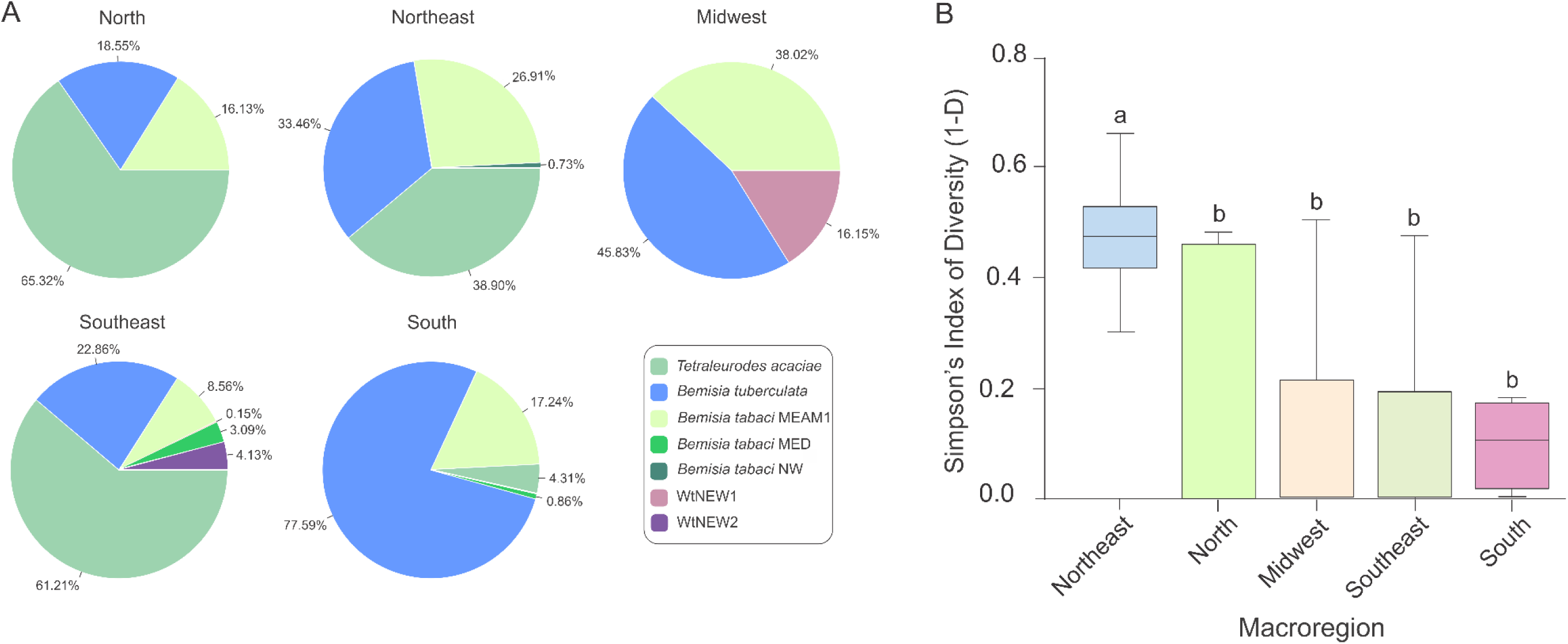
Composition and species diversity of whitefly populations differ among Brazilian regions. **A**. Pie charts represent the distribution of the 1,385 individuals genotyped in this study in the five geographic regions of Brazil. **B**. Boxplots correspond to Simpson’s index of diversity (1-D) calculated for each geographic region. The index was first calculated for each sampled site and grouped by geographic region. Different letters indicate significant differences between groups according to the non-parametric Kruskal-Wallis test followed by *post hoc* multiple comparison test (*p*<0.05).

### No begomoviruses detected infecting cassava

To verify the presence of begomoviruses infecting cassava, we analyzed leaves sampled in some of the fields where whiteflies were collected (Supp. Table S1). Based on PCR detection using universal primers for begomoviruses, all plants were negative.

## Discussion

Vectors play an essential role during the life cycle of plant viruses, directly affecting their ecology and evolution (74-76). Usually, a group of plant viruses establishes a very specific interaction with only one or a few related species of vectors, making virus ecology strongly dependent on that of its vector (74). It has been suggested that the natural host range of a virus is dependent on its vector’s host range, as most plant viruses have greater specificity for the vector than for the plant host (77, 78). Indeed, the existence of a competent vector for transmission and able to colonize potential reservoir and recipient new hosts is a primary ecological factor driving host range expansion of viruses. Thus, vectors play an essential role during viral disease emergence and epidemics (12, 18, 77, 79). Understanding ecological factors, such as vector species dynamics in crops, might provide important clues about historical and current events of emergence or re-emergence of viral diseases, and even anticipate the potential for new ones to occur (80).

Although it could be suggested that there are no begomoviruses capable of infecting cassava in the Americas, the high diversity of begomoviruses reported in a broad range of cultivated and non-cultivated plants in several botanical families, including the Euphorbiaceae, make this highly unlikely (33, 81-89). Thus, the absence of a competent vector able to colonize cassava and transfer begomoviruses from wild plants to cassava, as previously suggested (47), seems to be a more plausible hypothesis to explain the lack of begomovirus epidemics in this crop.

Our country-wide survey of whiteflies associated with cassava in Brazil uncovered a high degree of species diversity and showed that *T. acaciae* and *B. tuberculata* are the prevalent species across the country. Non-*B. tabaci* species, including *B. tuberculata*, have been shown to be prevalent also in Colombia (48). In contrast, in Africa, endemic species of the *B. tabaci* complex are prevalent in cassava (80, 90, 91). Previous studies surveying whitefly diversity in South American countries failed to detect *T. acaciae* and *B. tuberculata* in crops other than cassava, indicating a very narrow host range, which may in fact be restricted to cassava or at least to cultivated plants (51, 54, 92).

*Bt*MEAM1 and *Bt*NW are reported here for the first time in cassava in Brazil. *Bt*MEAM1 was the third most prevalent species, representing 18% of the genotyped individuals, and with similar incidence to *T. acaciae* and *B. tuberculata*. The failure of previous studies to detect *Bt*MEAM1 in cassava may have been due to the small number of samples analyzed. The wide distribution and prevalence of *Bt*MEAM1 in the main agroecological zones in Brazil has been well established, mostly in association with annual crops such as soybean, cotton, common bean and tomato (54). In these crops, *Bt*MEAM1 has a great reproductive capacity, rapidly increasing its population. Interestingly, our data showed the higher prevalence of *Bt*MEAM1 to be in the Midwest, where extensive agriculture predominates. The harvest of annual crops in the Midwest might cause the migration of the insect to semiperennial hosts such as cassava, which could explain why in some sites where *Bt*MEAM1 predominated among adults, it was not detected as nymphs (e.g., sites MT5, MT6, PR4).

It will be important to monitor *Bt*MEAM1 populations in cassava over the next years, to assess its possible adaptation to this host. The fact that we collected *Bt*MEAM1 nymphs at several locations suggests that this process may already be under way. We also detected *Bt*MED, a worrying result given the recent introduction of this species in the Brazil and its potential to displace other species, including *Bt*MEAM1 (93-95). *Bt*MED has disseminated quickly in the country, mainly in association with ornamental plants in greenhouses (54). Even though we detect *Bt*MED associated to cassava, we cannot infer its potential to effectively colonize this host since only adults were collected. The third species detected is the indigenous *Bt*NW. Although *Bt*MEAM1 partially displaced *Bt*NW in Brazil, this species can still be sporadically detected, mostly in association with non-cultivated hosts (50, 51, 54). It has been recurrently detected in *Euphorbia heterophylla*, suggesting a potential to colonize other species in the family Euphorbiaceae. However, the very low frequence with which it was detected and the absence of nymphs indicate that *Bt*NW is poorly adapted to cassava.

The identification of two putative new species highlights the remarkable genetic diversity of whiteflies. Interestingly, one of the new species was collected in the state of Mato Grosso, which corresponds to the region considered to be the domestication center of cassava (2-5). Further studies are needed to explore plant biodiversity in this region (96, 97), which might reveal a similar diversity of whiteflies which may be specifically adaptated to non-cultivated plant species due to long term co-evolution. The close phylogenetic relationship of the new species with non-*B. tabaci* whiteflies suggests that they are not virus vectors.

Whitefly species richness in cassava is just starting to be assessed, and may be greater than reported here. Based on morphological characters, Alonso, Racca-Filho (98) reported the presence of *Aleurothrixus aepim* and *Trialeurodes manihoti* colonizing cassava in the state of Rio de Janeiro. Although we did not analyze samples from that region, the failure to detect these species in other states suggests a restricted occurrence. Moreover, morphological characters alone are not always sufficient to classify whiteflies at the species level, and additional studies using molecular tools are needed to assess these molecularly uncharacterized whiteflies species (99).

Host suitability has been shown to be an important factor influencing the competitive capacity among species of the *B. tabaci* complex (94, 95, 100). Watanabe, Bello (95) demonstrated that displacement capacity between two invasive *B. tabaci* species was dependent on host suitability. While *Bt*MEAM1 displaced *Bt*MED only on tomato, *Bt*MED displaced *Bt*MEAM1 on sweet pepper and common bean. Luan, Xu (100) demonstrated that even in a host plant poorly suitable for *Bt*MEAM1, it was able to displace an indigenous species challenger. These authors demonstrated that even though host suitability may affect the speed of displacement, it may not affect the direction, as *Bt*MEAM1 always won the challenge (100). Interestingly, two or more species occurring sympatrically were detected in 51% of the fields analyzed in our study. In sites where *Bt*MEAM1 and *B. tuberculata* co-occurred, *B. tuberculata* predominated, suggesting a higher competitive capacity. Nonetheless, in all other combinations of co-occurring species, no differences in prevalence were observed. Thus, competitive capacity is unlikely to explain the low prevalence of *Bt*MEAM1, or the differences observed between *T. acaciae* and *B. tuberculata*.

Host adaptation may be a more important component affecting the low predominance of *Bt*MEAM1 in cassava, as previously suggested (47). The inability of *Bt*MEAM1 and *Bt*MED to colonize domesticated cassava efficiently has been demonstrated under experimental conditions (45, 46, 102, 103). Carabali, Montoya-Lerma (45), evaluating the colonization potential of *Bt*MEAM1 in three commercial cassava genotypes, demonstrated that only in one of them did *Bt*MEAM1 complete its development cycle from eggs to adult, and even then, at very low rates (0.003%). Using an electrical penetration graph assay, Milenovic, Wosula (103) demonstrated the inability of *Bt*MED to feed in cassava plants. Adults of this species spent a very short time ingesting cassava phloem sap compared to sap from a suitable host, suggesting that they would die by starvation in the field. Furthermore, the low efficiency of whiteflies of the *Bt*MED mitochondrial subgroups Q1 and Q2 in using cassava as a host has also been demonstrated (102). Oviposition and adult survival rates were very low, and development from eggs to adults was not observed. Although these studies were conducted under experimental conditions, the low predominance of *Bt*MEAM1 and *Bt*MED shown here and in other field surveys in Africa (90, 91, 101) strongly indicates a low adaptation of these species to cassava.

Nevertheless, our results indicate an ongoing adaptation process of *Bt*MEAM1 to cassava, with the detection of nymphs and adults in the same field. Interestingly, Carabali, Bellotti (47) demonstrated a gradual increase in the rate of reproduction and development of *Bt*MEAM1 after successive passages on plants phylogenetically related to the genus *Manihot* (*Euphorbia pulcherrima* and *Jatropha gossypiifolia*), indicating the potential of this whitefly species to become adapted to cassava through intermediate hosts. Furthermore, successful reproduction in the wild relative *M. esculenta* ssp. *flabellifolia* indicates that this plant may constitute an intermediate host leading to adaptation (46). This plant has been reported to be widely spread in the Amazon basin and the Midwest region of Brazil (97). Interestingly, our data showed the higher prevalence of *Bt*MEAM1 to be in the Midwest. Although we cannot establish a cause and effect relationship, it is reasonable to speculate that *M. esculenta* ssp. *flabellifolia* could be acting as an intermediate host mediating adaptation. A survey addressing whitefly diversity in this host should be necessary to test this hypothesis.

In Brazil, cassava is predominantly grown as a subsistence crop, usually side by side with other vegetables and with a high incidence of weeds. Growing cassava in a heterogenous environment, especially in the presence of related plants, may increase the adaptation potential of *Bt*MEAM1 and other species of the complex such as *Bt*MED, which we also detected in the open field. A high diversity of plants in cassava fields may allow an overlapping of ecological niches for distinct whitefly species, which under enough selection pressure may gradually adapt to new hosts. The sympatric occurrence of *T. acaciae, B. tuberculata* and *Bt*MEAM1, supports the role of botanical heterogeneity in shaping the composition of whitefly populations associated with cassava. A similar pattern was observed in Colombia, with 66% of the surveyed sites showing at least two species occurring sympatrically (48). Moreover, a predominance of one species in a given developmental stage and a different one in another stage (e.g., nymphs *vs* adults) at the same site suggests that other hosts may sustain reproduction and development, with adults migrating to cassava.

*Euphorbia heterophylla* (family Euphorbiaceae) is an invasive weed widely spread across Brazil and associated with several crops (89, 104). The presence of *E. heterophylla* plants in association with cassava (Figure 1A) and the fact that it was the most suitable host for *Bt*MEAM1 in Brazil out seven tested (105) shows its potential to act as an intermediate host mediating *Bt*MEAM1 adaptation. *E. heterophylla* has been frequently associated with the begomovirus

*Euphorbia yellow mosaic virus* (EuYMV) (89). Barreto, Hallwass (106) demonstrated that this plant is also a host of *Tomato severe rugose virus* (ToSRV), which even at a very low titer was transmitted to tomato plants, demonstrating the potential of *E. heterophylla* to act as a reservoir host. Surprisingly, considering that *E. heterophylla* and tomato belong to distinct botanical families, EuYMV is able to infect tomato (106). The closer botanical relationship between *E. heterophylla* and cassava may indicate a higher potential of EuYMV to infect cassava. The presence of EuYMV-infected *E. heterophylla* in cassava fields, as observed in this study (Figure 1A), its suitability as a host for *Bt*MEAM1, and the high efficiency of EuYMV transmission by *Bt*MEAM1 (107), suggest that EuYMV may have spillover potential to cassava. Experiments are ongoing in our laboratory to assess this spillover potential.

The emergence of begomoviruses in tomato crops in Brazil followed the introduction of *Bt*MEAM1 (32, 33), demonstrating the role of vector populations in promoting viral host range expansion and consequently epidemics. Thus, the adaptation of whiteflies to cassava could facilitate the emergence of begomoviruses in this crop. The establishment of management strategies to prevent or at least delaying the adaptation process is therefore necessary. *Bemisia tabaci* species may disperse across long distances international trade routes (108). Thus, preventing the introduction of cassava-adapted *B. tabaci* species from Africa should also be a priority.

### Data accessibility

mtCOI sequences obtained in this study were deposited in GenBank under accession numbers: MT901081 to MT901172 and MT904381 to MT904382. For detailed information see Supplementary Table S2.

## Author’s contribution

CADX, FMZ and RKS contributed to the design and implementation of the study; CADX, AMN, VHB, LFMW, MAJ, LFB, JEABJ, AJB, RFC, ESG, JHJ, GK, GSAL, CAL, RNM, KFCP, FNS, RRS, ENS and JWPS collected whitefly and leaf samples; CADX, AMN, VHB and LFMW processed whitefly samples; CADX performed the analyses and drafted the manuscript; CADX and FMZ prepared the final version of the manuscript.

## Acknowledgements

This work was funded by CAPES (Financial code 01), CNPq (409599/2016-6) and Fapemig (APQ-03276-18) grants to FMZ.

