## Supplementary figure S1 for "Assessing whitefly diversity to infer about begomovirus dynamics in cassava in Brazil"

**Supplementary Table S1.** Number and locations of samples used for begomovirus detection.

| Sample* | Location, state | Number of plants analyzed | Begomovirus-like symptoms | PCR result |
|---|---|---|---|---|
| BA2 | LEM, BA | 1 | nd | negative |
| DF1 | Planaltina, DF | 1 | nd | negative |
| DF2 | Planaltina, DF | 1 | nd | negative |
| DF3 | Planaltina, DF | 1 | nd | negative |
| MG1 | Ouro Fino, MG | 10 | non-symptoms | negative |
| MG10 | Florestal, MG | 10 | non-symptoms | negative |
| MG12 | Divinópolis, MG | 5 | non-symptoms | negative |
| MG13 | Viçosa, MG | 10 | non-symptoms | negative |
| MG19 | Caparaó, MG | 8 | non-symptoms | negative |
| MG2 | Pouso Alegre, MG | 10 | non-symptoms | negative |
| MG3 | Careaçu, MG | 5 | non-symptoms | negative |
| MG4 | Lambari, MG | 5 | non-symptoms | negative |
| MG5 | Lima Duarte, MG | 5 | non-symptoms | negative |
| MG6 | RioPomba, MG | 5 | non-symptoms | negative |
| MG7 | Florestal, MG | 5 | non-symptoms | negative |
| MG8 | Florestal, MG | 8 | non-symptoms | negative |
| MG9 | Florestal, MG | 5 | non-symptoms | negative |
| MT1 | Canarana, MT | 10 | non-symptoms | negative |
| MT2 | Canarana, MT | 1 | non-symptoms | negative |
| MT4 | Pedra Preta, MT | 1 | nd | negative |
| MT5 | Pedra Preta, MT | 1 | nd | negative |
| MT6 | Pedra Preta, MT | 1 | nd | negative |
| PA1 | Brasil Novo, PA | 10 | non-symptoms | negative |
| PA2 | Vitória do Xingu, PA | 8 | non-symptoms | negative |
| PA3 | Altamira, PA | 8 | non-symptoms | negative |
| PA4 | Altamira, PA | 2 | non-symptoms | negative |
| PR4 | Sertanópolis, PR | 2 | nd | negative |
| SP11 | Oleo, SP | 2 | nd | negative |
| SP9 | Casa Branca, SP | 2 | nd | negative |
| **Total** | | 143 | | |

* For detailed information about samples see Table 1. AL, Alagoas; BA, Bahia; DF, Distrito Federal; ES, Espírito Santo; GO, Goiás; MG, Minas Gerais; MT, Mato Grosso; PA, Pará; PI, Piauí; PR, Paraná; SC, Santa Catarina; SP, São Paulo. nd: nondetermined.

**Supplementary Table S2.** Whiteflies isolates obtained in this study and GenBank access numbers are shown.

| **Sample**[1] | **Collection date** | **Country** | **Host** | **Specie** | **Isolate**[2] | **GenBank access #** |
|---|---|---|---|---|---|---|
| MG1 | May-18 | Brazil | *Manihot esculenta* | *Tetraleurodes acaciae* | AMG1-7 | MT901081 |
| MG1 | May-18 | Brazil | *Manihot esculenta* | *Tetraleurodes acaciae* | NMG1-1 | MT901082 |
| MG2 | Jul-18 | Brazil | *Manihot esculenta* | *Tetraleurodes acaciae* | AMG2-1 | MT901083 |
| MG2 | Jul-18 | Brazil | *Manihot esculenta* | *Tetraleurodes acaciae* | NMG2-1 | MT901084 |
| MG3 | Feb-18 | Brazil | *Manihot esculenta* | *Tetraleurodes acaciae* | AMG3-2 | MT901085 |
| MG3 | Feb-18 | Brazil | *Manihot esculenta* | *Tetraleurodes acaciae* | NMG3-3 | MT901086 |
| MG4 | Jun-18 | Brazil | *Manihot esculenta* | *Tetraleurodes acaciae* | AMG4-2 | MT901087 |
| MG4 | Jun-18 | Brazil | *Manihot esculenta* | *Tetraleurodes acaciae* | NMG4-2 | MT901088 |
| MG5 | Feb-18 | Brazil | *Manihot esculenta* | *Tetraleurodes acaciae* | AMG5-11 | MT901089 |
| MG5 | Feb-18 | Brazil | *Manihot esculenta* | *Tetraleurodes acaciae* | NMG5-1 | MT901090 |
| MG7 | Feb-18 | Brazil | *Manihot esculenta* | *Tetraleurodes acaciae* | AMG7-1 | MT901091 |
| MG7 | Feb-18 | Brazil | *Manihot esculenta* | *Tetraleurodes acaciae* | NMG7-1 | MT901092 |
| MG8 | Feb-18 | Brazil | *Manihot esculenta* | *Tetraleurodes acaciae* | AMG8-1 | MT901093 |
| MG9 | Mar-18 | Brazil | *Manihot esculenta* | *Tetraleurodes acaciae* | AMG9-1 | MT901094 |
| MG9 | Mar-18 | Brazil | *Manihot esculenta* | *Tetraleurodes acaciae* | NMG9-1 | MT901095 |
| MG10 | Feb-18 | Brazil | *Manihot esculenta* | *Tetraleurodes acaciae* | AMG10-1 | MT901096 |
| MG10 | Feb-18 | Brazil | *Manihot esculenta* | *Tetraleurodes acaciae* | NMG10-1 | MT901097 |
| MG11 | Mar-18 | Brazil | *Manihot esculenta* | *Tetraleurodes acaciae* | AMG11-1 | MT901098 |
| MG11 | Mar-18 | Brazil | *Manihot esculenta* | *Tetraleurodes acaciae* | AMG11-8 | MT901099 |
| MG11 | Mar-18 | Brazil | *Manihot esculenta* | *Tetraleurodes acaciae* | NMG11-1 | MT901100 |
| MG12 | May-18 | Brazil | *Manihot esculenta* | *Tetraleurodes acaciae* | AMG12-7 | MT901101 |
| MG12 | May-18 | Brazil | *Manihot esculenta* | *Tetraleurodes acaciae* | NMG12-1 | MT901102 |
| MG14 | Mar-18 | Brazil | *Manihot esculenta* | *Tetraleurodes acaciae* | AMG14-5 | MT901103 |
| MG15 | Mar-18 | Brazil | *Manihot esculenta* | *Tetraleurodes acaciae* | AMG15-1 | MT901104 |
| MG15 | Mar-18 | Brazil | *Manihot esculenta* | *Tetraleurodes acaciae* | NMG15-2 | MT901105 |
| MG16 | Mar-18 | Brazil | *Manihot esculenta* | *Tetraleurodes acaciae* | AMG16-1 | MT901106 |
| MG17 | Mar-18 | Brazil | *Manihot esculenta* | *Tetraleurodes acaciae* | AMG17-9 | MT901107 |
| MG18 | Mar-18 | Brazil | *Manihot esculenta* | *Tetraleurodes acaciae* | AMG18-5 | MT901108 |
| MG19 | Feb-18 | Brazil | *Manihot esculenta* | *Tetraleurodes acaciae* | AMG19-10 | MT901109 |
| MG19 | Feb-18 | Brazil | *Manihot esculenta* | *Tetraleurodes acaciae* | NMG19-4 | MT901110 |
| ES1 | Jan-18 | Brazil | *Manihot esculenta* | *Tetraleurodes acaciae* | AES1-11 | MT901111 |
| ES1 | Jan-18 | Brazil | *Manihot esculenta* | *Tetraleurodes acaciae* | NES1-1 | MT901112 |
| ES2 | Jan-18 | Brazil | *Manihot esculenta* | *Tetraleurodes acaciae* | AES2-2 | MT901113 |
| ES2 | Jan-18 | Brazil | *Manihot esculenta* | *Tetraleurodes acaciae* | NES2-1 | MT901114 |
| ES2 | Jan-18 | Brazil | *Manihot esculenta* | *Tetraleurodes acaciae* | NES2-12 | MT901115 |
| ES3 | Jan-18 | Brazil | *Manihot esculenta* | *Tetraleurodes acaciae* | AES3-2 | MT901116 |
| ES3 | Jan-18 | Brazil | *Manihot esculenta* | *Tetraleurodes acaciae* | NES3-1 | MT901117 |
| PA1 | Jan-18 | Brazil | *Manihot esculenta* | *Tetraleurodes acaciae* | APA1-1 | MT901118 |
| PA1 | Jan-18 | Brazil | *Manihot esculenta* | *Tetraleurodes acaciae* | APA1-11 | MT901119 |
| PA1 | Jan-18 | Brazil | *Manihot esculenta* | *Tetraleurodes acaciae* | NPA1-11 | MT901120 |

| | | | | | | |
|---|---|---|---|---|---|---|
| PA2 | Jan-18 | Brazil | *Manihot esculenta* | *Tetraleurodes acaciae* | APA2-16 | MT901121 |
| PA2 | Jan-18 | Brazil | *Manihot esculenta* | *Tetraleurodes acaciae* | NPA2-2 | MT901122 |
| PA3 | Aug-18 | Brazil | *Manihot esculenta* | *Tetraleurodes acaciae* | APA3-6 | MT901123 |
| PA3 | Aug-18 | Brazil | *Manihot esculenta* | *Tetraleurodes acaciae* | NPA3-2 | MT901124 |
| PA4 | Jan-18 | Brazil | *Manihot esculenta* | *Tetraleurodes acaciae* | NPA4-16 | MT901125 |
| AL1 | Jul-18 | Brazil | *Manihot esculenta* | *Tetraleurodes acaciae* | NAL1-17 | MT901126 |
| AL2 | Apr-18 | Brazil | *Manihot esculenta* | *Tetraleurodes acaciae* | AAL2-10 | MT901127 |
| AL3 | Apr-18 | Brazil | *Manihot esculenta* | *Tetraleurodes acaciae* | AAL3-6 | MT901128 |
| AL4 | Apr-18 | Brazil | *Manihot esculenta* | *Tetraleurodes acaciae* | AAL4-2 | MT901129 |
| AL4 | Apr-18 | Brazil | *Manihot esculenta* | *Tetraleurodes acaciae* | NAL4-7 | MT901130 |
| AL5 | Apr-18 | Brazil | *Manihot esculenta* | *Tetraleurodes acaciae* | NAL5-4 | MT901131 |
| BA1 | Dec-17 | Brazil | *Manihot esculenta* | *Tetraleurodes acaciae* | ABA1-8 | MT901132 |
| PI1 | Apr-18 | Brazil | *Manihot esculenta* | *Tetraleurodes acaciae* | NPI1-19 | MT901133 |
| BA1 | Jul-18 | Brazil | *Manihot esculenta* | *Bemisia tuberculata* | AMG2-9 | MT901134 |
| MG3 | Feb-18 | Brazil | *Manihot esculenta* | *Bemisia tuberculata* | AMG3-1 | MT901135 |
| MG3 | Feb-18 | Brazil | *Manihot esculenta* | *Bemisia tuberculata* | NMG3-1 | MT901136 |
| MG6 | Apr-18 | Brazil | *Manihot esculenta* | *Bemisia tuberculata* | AMG6-6 | MT901137 |
| MG10 | Feb-18 | Brazil | *Manihot esculenta* | *Bemisia tuberculata* | AMG10-7 | MT901138 |
| MG10 | Feb-18 | Brazil | *Manihot esculenta* | *Bemisia tuberculata* | NMG10-12 | MT901139 |
| MG12 | May-18 | Brazil | *Manihot esculenta* | *Bemisia tuberculata* | AMG12-3 | MT901140 |
| MG13 | Aug-18 | Brazil | *Manihot esculenta* | *Bemisia tuberculata* | AMG13-3 | MT901141 |
| MG15 | Mar-18 | Brazil | *Manihot esculenta* | *Bemisia tuberculata* | NMG15-1 | MT901142 |
| MG16 | Mar-18 | Brazil | *Manihot esculenta* | *Bemisia tuberculata* | AMG16-4 | MT901143 |
| MG18 | Mar-18 | Brazil | *Manihot esculenta* | *Bemisia tuberculata* | AMG18-18 | MT901144 |
| MT1 | Dec-17 | Brazil | *Manihot esculenta* | *Bemisia tuberculata* | AMT1-16 | MT901145 |
| MT1 | Dec-17 | Brazil | *Manihot esculenta* | *Bemisia tuberculata* | NMT1-16 | MT901146 |
| MT2 | Dec-17 | Brazil | *Manihot esculenta* | *Bemisia tuberculata* | NMT2-4 | MT901147 |
| ES3 | Jan-18 | Brazil | *Manihot esculenta* | *Bemisia tuberculata* | AES3-4 | MT901148 |
| ES3 | Jan-18 | Brazil | *Manihot esculenta* | *Bemisia tuberculata* | NES3-5 | MT901149 |
| PA3 | Aug-18 | Brazil | *Manihot esculenta* | *Bemisia tuberculata* | APA3-7 | MT901150 |
| PA4 | Jan-18 | Brazil | *Manihot esculenta* | *Bemisia tuberculata* | APA4-1 | MT901151 |
| PR1 | Mar-18 | Brazil | *Manihot esculenta* | *Bemisia tuberculata* | APR1-16 | MT901152 |
| PR2 | Mar-18 | Brazil | *Manihot esculenta* | *Bemisia tuberculata* | APR2-18 | MT901153 |
| GO2 | Mar-18 | Brazil | *Manihot esculenta* | *Bemisia tuberculata* | AGO2-16 | MT901154 |
| AL1 | Jul-18 | Brazil | *Manihot esculenta* | *Bemisia tuberculata* | NAL1-2 | MT901155 |
| AL2 | Apr-18 | Brazil | *Manihot esculenta* | *Bemisia tuberculata* | AAL2-1 | MT901156 |
| AL2 | Apr-18 | Brazil | *Manihot esculenta* | *Bemisia tuberculata* | NAL2-1 | MT901157 |
| AL3 | Apr-18 | Brazil | *Manihot esculenta* | *Bemisia tuberculata* | AAL3-3 | MT901158 |
| AL3 | Apr-18 | Brazil | *Manihot esculenta* | *Bemisia tuberculata* | NAL3-1 | MT901159 |
| AL4 | Apr-18 | Brazil | *Manihot esculenta* | *Bemisia tuberculata* | AAL4-4 | MT901160 |
| AL4 | Apr-18 | Brazil | *Manihot esculenta* | *Bemisia tuberculata* | NAL4-1 | MT901161 |
| BA1 | Dec-17 | Brazil | *Manihot esculenta* | *Bemisia tuberculata* | ABA1-12 | MT901162 |
| PI1 | Apr-18 | Brazil | *Manihot esculenta* | *Bemisia tuberculata* | NPI1-16 | MT901163 |
| GO1 | Mar-18 | Brazil | *Manihot esculenta* | *Bemisia tabaci* MEAM1 | NGO1-1 | MT901164 |
| GO2 | Mar-18 | Brazil | *Manihot esculenta* | *Bemisia tabaci* MEAM1 | AGO2-2 | MT901165 |

| | | | | | | |
|---|---|---|---|---|---|---|
| GO2 | Mar-18 | Brazil | *Manihot esculenta* | *Bemisia tabaci* MEAM1 | NGO2-5 | MT901166 |
| AL2 | Apr-18 | Brazil | *Manihot esculenta* | *Bemisia tabaci* MEAM1 | AAL2-7 | MT901167 |
| AL3 | Apr-18 | Brazil | *Manihot esculenta* | *Bemisia tabaci* MEAM1 | AAL3-8 | MT901168 |
| AL5 | Apr-18 | Brazil | *Manihot esculenta* | *Bemisia tabaci* MEAM1 | AAL5-3 | MT901169 |
| BA1 | Dec-17 | Brazil | *Manihot esculenta* | *Bemisia tabaci* MEAM1 | ABA1-20 | MT901170 |
| AL2 | Apr-18 | Brazil | *Manihot esculenta* | *Bemisia tabaci* NW | AAL2-11 | MT901171 |
| MT1 | Dec-17 | Brazil | *Manihot esculenta* | *Tetraleorodes sp.*[3] | AMT1-16 | MT901172 |
| MG6 | Apr-18 | Brazil | *Manihot esculenta* | *Bemisia sp.*[4] | AMG6-16 | MT904381 |
| MG6 | Apr-18 | Brazil | *Manihot esculenta* | *Bemisia sp.*[4] | NMG6-4 | MT904382 |

[1] For detailed information about samples see Table 1.

[2] First letter of isolates names indicates if a given sequence came from adults (A) or nymphs (N), followed for two letters representing the state from where it was collected: AL, Alagoas; BA, Bahia; DF, Distrito Federal; ES, Espírito Santo; GO, Goiás; MG, Minas Gerais; MT, Mato Grosso; PA, Pará; PI, Piauí; PR, Paraná; SC, Santa Catarina; SP, São Paulo.

[3] Correspond to WtNEW1.

[4] Correspond to WtNEW2.

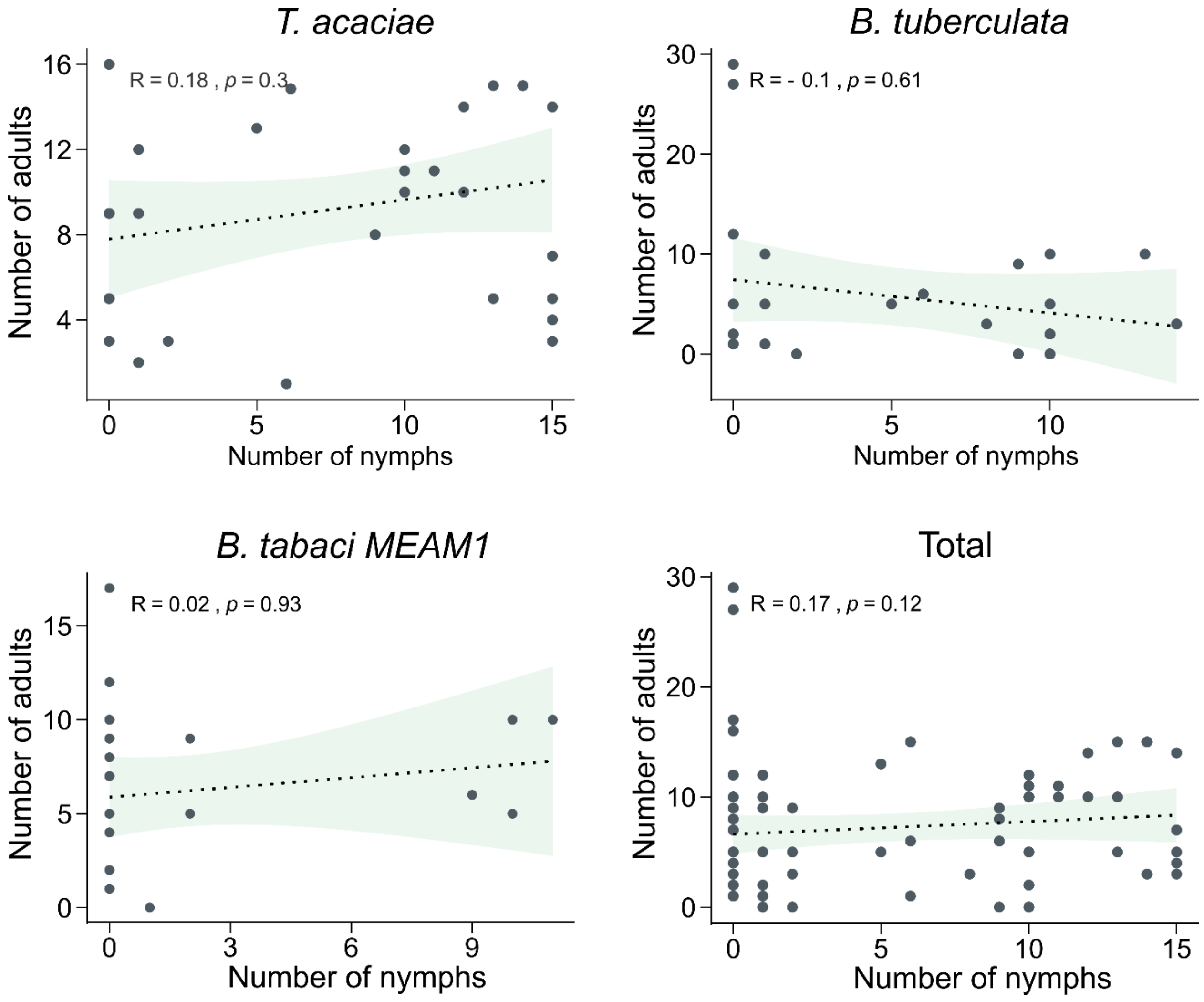


**Supplementary Figure S1**. Spearman's rank correlation coefficient analysis comparing number of nymphs and adults for the three more abundant whiteflies specie. Each dot represent a sampled field. Only fields where nymphs and adults or where only one phase was observed were included in this analysis. Scatter plot showing 95% of confidence interval (light green) are shown. Total, correspond all three specie ploted together.
